# MOSAIC: Model-based, Subgroup-Aware Identification of Driver Mutations in Cancer

**DOI:** 10.64898/2026.04.29.721672

**Authors:** Kiersten Campbell, Matthew A. Reyna

## Abstract

In cancer genomics, recurrent patterns of mutual exclusivity within a gene set can indicate shared biological context and involvement in tumorigenesis. However, existing methods are not designed to distinguish between mutual exclusivity arising from meaningful biological interactions from those influenced by heterogeneity between underlying patient subpopulations. In this work, we introduce MOSAIC, a novel statistical framework that models patient subgroup heterogeneity in mutual exclusivity analyses. In experiments with simulated data and real data from The Cancer Genome Atlas, we show that MOSAIC amplifies subgroup-specific mutual exclusivity signals, including between *IDH1* and *IDH2* in young low grade glioma patients, while reducing the effect of signals produced by underlying subgroup structures, such as distinct genomic lineages associated with histological subtypes of endometrial cancer. Finally, we demonstrate that MOSAIC is more powerful than existing *p*-value combination methods for patient subgroup stratification. MOSAIC is available as an open-source tool at https://github.com/reynalab/mosaic.

## 1 Introduction

Extensive genomic heterogeneity in cancer challenges the identification and interpretation of cancer driver genes that cause tumor initiation and progression. Many mutations are found in less than 5-10% of tumors within the same tissue type, which can frustrate both our biological knowledge and clinical interventions, even in large tumor cohorts [11,16,36,3]. To better address intertumor heterogeneity for the identification of novel cancer driver genes, a class of computational methods evaluates recurrent patterns of mutations within gene sets, rather than evaluating recurrent mutations in individual genes, thereby leveraging the heterogeneous patterns of mutations in tumor cohorts to identify driver genes. Mutually exclusive mutations, or patterns of mutations in a given gene set that rarely co-occur in the same tumor, is a common pattern observed in known driver genes (e.g., *KRAS* and *BRAF* are recurrently mutated driver genes in colorectal cancers that are rarely altered in the same patient [27]). Several methods have been developed to identify mutually exclusive gene sets *de novo* from genomics data to identify novel interactions and rare driver genes [28].

Some *de novo* mutual exclusivity methods quantify the significance of an observed pattern of mutual exclusivity through combinatorial scores, including Dendrix [35] and Multi-Dendrix [17]. However, combinatorial scores can be confounded by the mutation rates of the genes in a given gene set. For example, *TTN*, a long gene that naturally accumulates many passenger mutations, is not a cancer driver gene, but it often presents as a false positive in pairs with infrequently mutated genes by chance [16]. To address this issue, statistical models, including WeSME [14], MEGSA [12], CoMEt [19], WeXT [18], and DIALECT [30], quantify the observed number of mutually exclusive mutations in a gene set relative to a null distribution of mutual exclusivity patterns generated by chance alone.

Although mutual exclusivity can be a useful signal for identifying evidence of selection in cancer, there are many causes of mutual exclusivity in cancer genomes that may or may not implicate genes as driver genes. For example, mutual exclusivity may indicate shared biological context (e.g., participation in the same pathway) that is altered in cancer, synthetic lethality, or distinct genomic lineages within a cancer type, but these different causes have different interpretations [6,5,8]. Existing work also demonstrates that the prevalences of known cancer driver mutations vary across patient subpopulations, which may influence patterns of mutual exclusivity. For example, many known driver genes, including *TP53,AKT1*, and *GATA3*, occur at significantly different rates in older (65+ years old) and younger (18-64 years old) breast cancer patients [1, 13, 37]. Similarly, *TP53, CDKN2A*, and *TERT* mutations were found to be enriched in metastatic tumors relative to localized tumors [20]. These differences may contribute to a pattern of mutually exclusive mutations in a heterogeneous patient cohort even when they occur in genes that are not cancer driver genes.

Indeed, the sources of a pattern of mutual exclusivity between genes affect our interpretation of the genes. Despite that, existing mutual exclusivity methods may address intertumor heterogeneity at the resolution of individual patients, but they largely do not explicitly control for variation between groups of patients within a single tissue type. As a result, it can be difficult to determine the degree to which the genes identified by these methods reflect biological interactions that are indicative of cancer drivers or are driven by underlying patient subgroup structures within a tumor cohort.

To the best of our knowledge, no existing mutual exclusivity method explicitly addresses heterogeneity with patient subgroups. CoMEt [19] optionally considers cancer subtypes, but its approach may not account for mutual exclusivity specific to one or more patient subgroups, and it has not been tested on other subgroup definitions; see Section 3.1 for a deeper analysis. Therefore, an approach that resolves intertumor heterogeneity and subgroup heterogeneity in a robust statistical model is needed to better contextualize mutual exclusivity signals.

Here, we introduce MOSAIC, a <u>mo</u>del-based, subgroup-<u>a</u>ware method for the identification of driver mutations in <u>c</u>ancer. MOSAIC builds upon existing *de novo* mutual exclusivity methods by modeling mutational heterogeneity between user-defined patient subgroups. As a result, MOSAIC:

1. contextualizes mutual exclusivity signals that arise primarily through differences in gene mutation rates between subgroups, including those associated with subtype-specific genomic lineages;
2. amplifies mutual exclusivity signals specific to one or more subgroups;
3. retains the statistical power of subgroup-unaware approaches in the absence of subgroup-related heterogeneity.

We tested MOSAIC on both simulated and real genomics data. In particular, in this work, we demonstrate MOSAIC’s ability to model subgroup-specific genomic lineages within histological subtypes of endometrial cancer and to amplify signals of mutual exclusivity specific to younger low-grade glioma patients from The Cancer Genome Atlas Pan-Cancer Atlas Project [32].

## 2 Methods

### 2.1 Definitions

Let *A* = [*a*_*ij*_] ∈ {0, 1} ^*m ×n*^ be an observed binary matrix representing the occurrence of *m* events across *n* samples or patients, where *a*_*ij*_ indicates the presence (*a*_*ij*_ = 1) or absence (*a*_*ij*_ = 0) of an event *i* in sample *j*. For the simplicity of presentation, we will most often describe MOSAIC in terms of genetic mutations in patient samples (i.e., *a*_*ij*_ =1 indicates that gene *i* is mutated in sample *j*), but MOSAIC can be used with other, more general binary matrices.

Two events *i* and *ℓ* are *mutually exclusive* in sample *j* if and only if one and only one of events *i* or *ℓ* is present in sample *j* (i.e., *a*_*ij*_ + *a*_*ℓj*_ =1 with either *a*_*ij*_ =1 and *a*_*ℓj*_ =0 *or a*_*ij*_ =0 and *a*_*ℓj*_ = 1). The number of mutually exclusive observations between events *i* and *ℓ* in *A* is

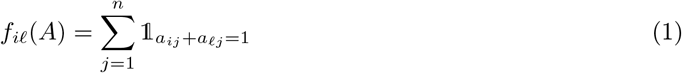

and the frequency of event *i* in *A* is

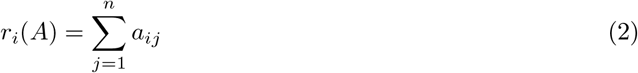

with **r**(*A*)= (*r*_1_(*A*),. .., *r*_*m*_(*A*)) denoting a sequence of event frequencies across *A*.

### 2.2 Subgroup-Unaware Mutual Exclusivity Test

The subgroup-unaware mutual exclusivity test assesses observed patterns of mutual exclusivity without considering patient subgroups.

We define

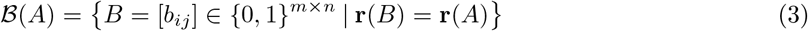

as the set of all possible binary mutation matrices with the same gene mutation rates observed in *A*.

Under the null hypothesis that the events *i* and *ℓ* are independent, conditioned on the mutation frequency of the events, we compute

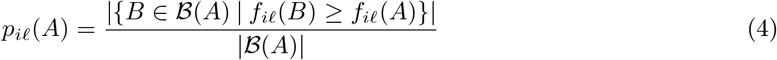

as the probability of observing as many or more mutually exclusive mutations between events *i* and *ℓ*.

Enumerating ℬ(*A*) is computationally intractable for large matrices because 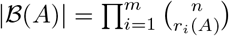. However, the exclusivity and co-occurence of mutations between two genes can be summarized by a 2 × 2 contingency table whose margins preserve the genes’ mutation rates. Indeed, the subgroup-unaware mutual exclusivity test in (4) for a pair of genes is equivalent to a one-sided Fisher’s exact test [9] on the contingency table, and is equivalent to special cases of CoMEt [19] and WExT [18]. Therefore, (4) can be computed more efficiently using the hypergeometric distribution

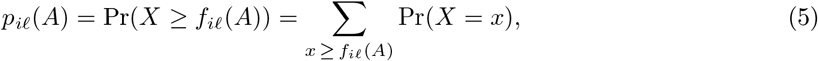

where *X* is a random variable, defined in terms of the hypergeometric distribution, corresponding to the number of mutually exclusive mutations under the null distribution in (3).

### 2.3 MOSAIC: A Subgroup-Aware Mutual Exclusivity Test

The subgroup-aware mutual exclusivity test, summarized in Figure 1, assesses observed patterns of mutual exclusivity while considering pre-defined patient subgroups.

**Fig. 1:**
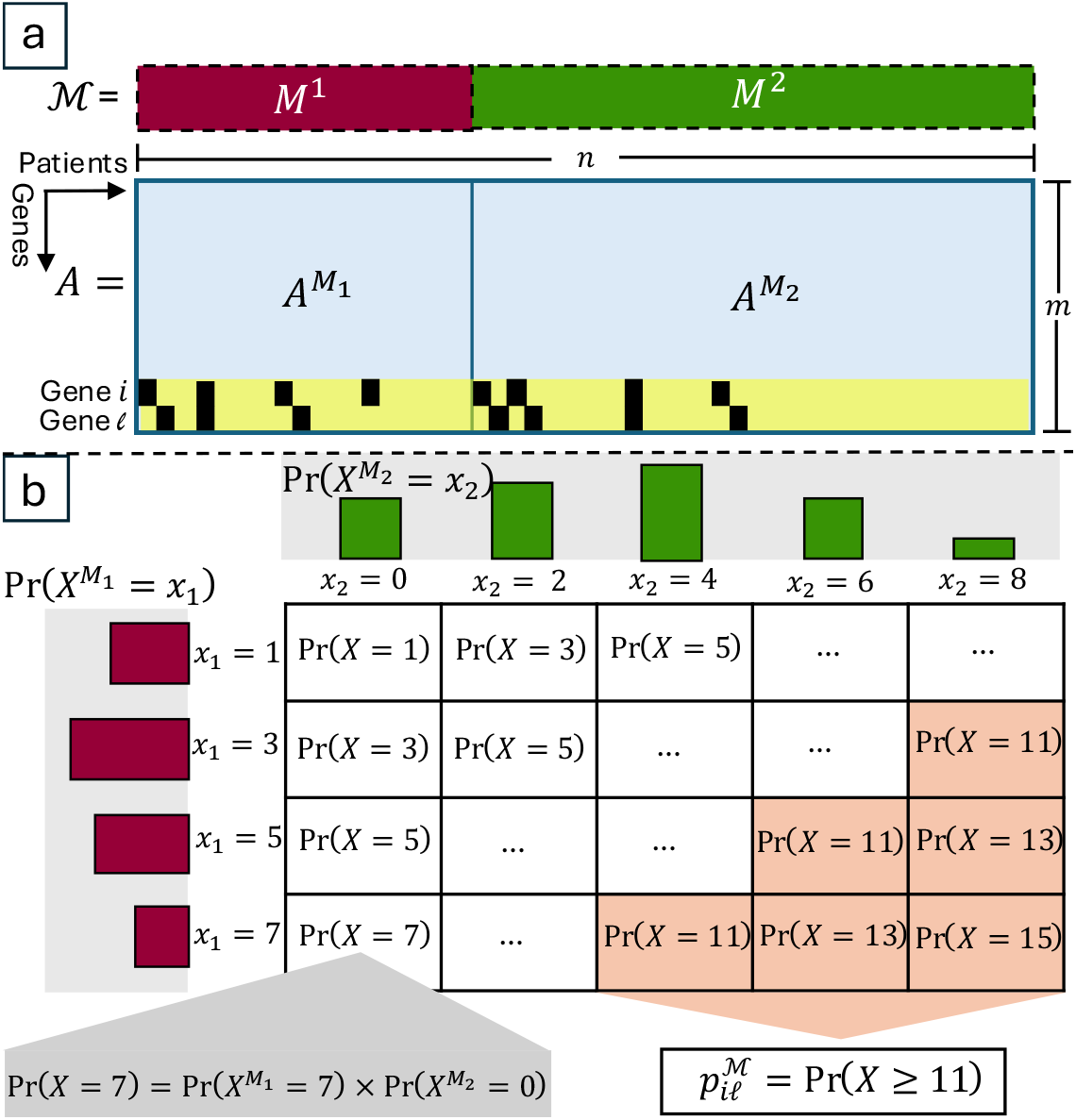
MOSAIC: a subgroup-aware mutual exclusivity framework. (a) Genomics data is represented in an example binary matrix, where *a*_*ij*_ = 1 (black bars) indicates an observed mutation in gene *i* in patient *j*. For simplicity, only mutations in two genes are shown. MOSAIC contextualizes an observed signal of mutual exclusivity, *f*_*iℓ*_(*A*) = 11, within patient subgroups, specified by *M* ^1^ and *M* ^2^. (b) This subgroup-aware approach generates a unified null distribution based on the possible unique configurations of each subgroup.

Let ℳ = {*M*_1_,. .., *M*_*k*_} be a partition of the patients or samples {1,.. ., *n*} into *k* pairwise distinct subgroups (i.e., 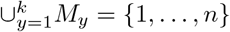 and *M*_*y*_ ∩ *M*_*z*_ = ∅ for every *y* ≠ *z*). We define *A*^*M*^ as the submatrix of *A* induced by *M* ∈ ℳ (i.e., the columns of *A* corresponding to the samples in *M*). We define

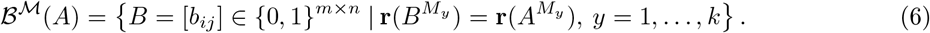

as the set of all possible binary mutation matrices with the same gene mutation rates observed in *A*^*M*^ for each *M* ∈ ℳ.

As in Section 2.2, enumerating ℬ^*ℳ*^(*A*) is computationally intractable for large matrices, but we can again formulate the subgroup-aware method as a generalization of Fisher’s exact test. The subgroup-aware null model preserves the mutation rates observed in each submatrix. This is equivalent to preserving the marginals of the 2 × 2 contingency table summarizing *A*^*M*^ for each *M* ∈ ℳ. Let 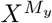 denote a random variable corresponding to the number of mutually exclusive samples in subgroup *M*_*y*_ under the null model for 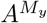, with its distribution given by the corresponding hypergeometric distribution. Since the subgroups are modeled independently, the probability of a mutual exclusivity pattern in the full cohort is the joint probability of the subgroup-specific events:

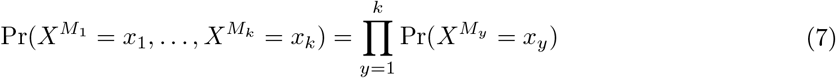

Multiple unique configurations of submatrices can produce the same number of mutually exclusive mutations in the overall cohort, so the probability of a given number of mutually exclusive mutations in the overall cohort is computed over all of the possible configurations of submatrices producing that signal in the full cohort:

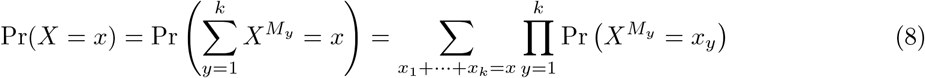

Note that the number of possible configurations of the sample subgroups, (*x*_1_, *x*_2_, …, *x*_*k*_), resulting in a given signal in the overall cohort, *x*, grows rapidly as the number of subgroups increases. Therefore, we restrict the computation to configurations in which each *x*_*y*_ has non-zero probability under the corresponding subgroup-specific hypergeometric distribution (i.e., the support), which considerably reduces the number of possible configurations in practice.

Therefore, the *p*-value for MOSAIC, the subgroup-aware mutual exclusivity test, is the sum of the probabilities in (8) that achieve as many or more mutually exclusive mutations as the observed mutation matrix:

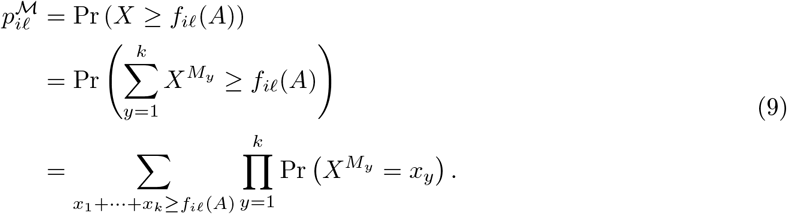

We note that our subgroup-aware approach, MOSAIC, preserves the mutual exclusivity test statistic *f*_*iℓ*_(*A*) of the subgroup-unaware approach but constrains the null model ℬ^ℳ^ from the subgroup-unaware approach so that ℬ^ℳ^(*A*) ⊆ ℬ(*A*). In particular, if we were to choose *k* =1 so that ℳ = {{1,. .., *n*}}, then ℬ^ℳ^(*A*) = ℬ(*A*) and the subgroup-aware approach reduces to the subgroup-unaware approach with a comparable *p*-value. Conversely, if we were to choose *k* = *n* so that ℳ = {{1},…, {*n*}}, then ℬ^ℳ^(*A*)= {*A*} and the subgroup-aware approach reduces to the trivial case with a *p*-value of 1.

### 2.4 Contextualization of Mutual Exclusivity Signals

In practice, users should run both subgroup-unaware (i.e., ℳ = {{1,.. ., *n*}}) and subgroup-aware (i.e., ℳ defined according to a sample covariate) approaches. Comparing the resulting *p*-values from both approaches helps to contextualize the observed signal of mutual exclusivity for a given gene pair. In these comparisons, users should consider the relative difference in *p*-values (e.g., differences of at least one order of magnitude) as indicative of meaningful changes in significance between approaches. Three cases can arise:

- 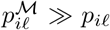: The mutual exclusivity signal appears to be less significant under the subgroup-aware model, indicating that the pattern of mutual exclusivity may reflect subgroup heterogeneity rather than underlying biological interactions between the genes.
- 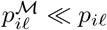: The mutual exclusivity signal appears to be more significant under the subgroup-aware model, indicating that the pattern of mutual exclusivity may be specific to one or more subgroups and was obscured in the overall cohort.
- 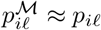: The mutual exclusivity signal is not significantly influenced by subgroup heterogeneity.

As with other statistical hypothesis tests, the *p*-values from (5) and (9) may be compared to a pre-defined threshold (e.g., *α* = 0.05) to determine the statistical significance of the observed signal of mutual exclusivity. Multiple testing corrections are typically needed when multiple pairs of genes are evaluated (e.g., [7, 23, 2]).

### 2.5 Implementation

We implemented MOSAIC in Python. We released our code, documentation, examples, experiments, and commands for reproducing the results in this manuscript under an open-source license on https://github.com/reynalab/mosaic.

## 3 Results

### 3.1 Example Data

To illustrate MOSAIC’s behavior on data with different sources of mutual exclusivity, we created an example mutation matrix with 6 genes, 12 patients, and 2 patient subgroups (see Figure 2).

**Fig. 2:**
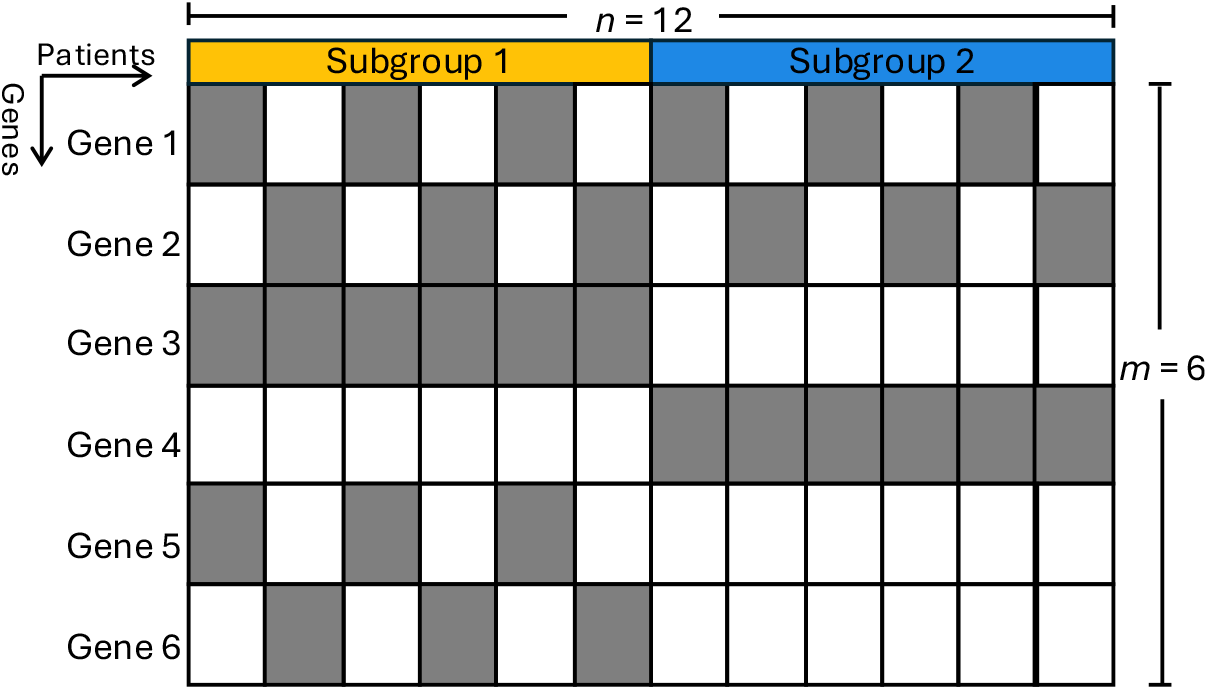
Toy data. An example mutation matrix illustrating different sources of mutual exclusivity between Genes 1 and 2, Genes 3 and 4, and Genes 5 and 6. Gray cells indicate that *a*_*ij*_ = 1, i.e., gene *i* is mutated in patient *j*.

First, Genes 1 and 2 have mutually exclusive mutations that are independent of the patient subgroups, illustrating genes with shared biological context. MOSAIC (9) and Fisher’s exact test (5) return similar *p*-values (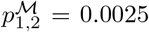 and *p*_1,2_ = 0.0011, respectively). MOSAIC returns a slightly larger *p*-value because of patient stratification of a small number of samples in the toy dataset, but both results are statistically significant results at a significance level of *α* = 0.05. Second, Genes 3 and 4 have mutually exclusive mutations that depend on the patient subgroups, illustrating mutation rate differences between the subgroups. MOSAIC returns a larger *p*-value than Fisher’s exact test (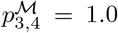 and *p*_3,4_ = 0.0011, respectively), demonstrating that MOSAIC reduces the effect of signals arising from underlying subgroup structure. Third, Genes 5 and 6 have mutually exclusive mutations only in one patient subgroup. MOSAIC returns a much smaller *p*-value than Fisher’s exact test (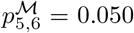 and *p*_5,6_ = 0.38, respectively), highlighting MOSAIC’s ability to localize subgroup-specific signals that are obscured in pooled analyses.

In comparison, CoMEt [17] can consider cancer subtypes by adding subtype indicator rows to the observed mutation matrix. Indeed, for a gene pair driven by subgroup structure (e.g., Genes 3 and 4), CoMEt produces a larger *p*-value with the subtype rows (*p* = 0.99 with the subtype indicator rows and *p* = 0.0005 without them, respectively). However, when a mutual exclusivity pattern is independent of the subgroups (e.g., Genes 1 and 2), CoMEt still returns the same *p*-values (*p* = 0.99 with the subtype rows and *p* = 0.0005 without them) as Genes 3 and 4 with subgroup-dependent exclusivity, treating different sources of heterogeneity equivalently. MOSAIC, on the other hand, reduces the effect of mutual exclusivity signals driven by subtype-specific lineages (Genes 3 and 4; 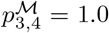) while preserving the significance of mutual exclusivity signals shared across subgroups (Genes 1 and 2; 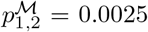). We also evaluated CoMEt’s behavior in cases where the mutual exclusivity signal is found in only one subgroup (e.g., Genes 5 and 6) with no mutations in a second subgroup. When the subgroup without mutations in Genes 5 and 6 is excluded from the matrix, CoMEt generates a smaller *p*-value than when this subgroup is included (*p* = 0.025 and *p* = 0.18, respectively). MOSAIC more strongly amplifies mutual exclusivity signals found in only one subgroup 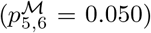 and maintains the same *p*-value whether this subgroup is included or excluded, intuitively localizing the subgroup-specific signal.

Finally, we compared MOSAIC to the meta-analysis methods of Fisher [9], Tippett [33], Pearson [26], Stouffer [31], and Mudholkar and George [24] for combining the *p*-values from Fisher’s exact test on each patient subgroup (Appendix Table 1). For Genes 1 and 2, MOSAIC returns a smaller *p*-value than Fisher’s method (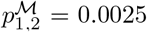 and *p*_1,2_ = 0.0175, respectively) and the other methods. For this pair, which exhibits a signal of mutual exclusivity that is independent of the patient subgroups, MOSAIC is a less powerful test than applying a subgroup-unaware approach, like Fisher’s exact test, to the entire sample cohort, but it is more powerful than applying *post-hoc p*-value combination approaches. For Gene 3 and 4, MOSAIC and the *p*-value combination methods perform identically (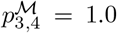 and *p*_3,4_ = 1.0, respectively) because the mutual exclusivity signal is perfectly confounded by subgroup structure. For Genes 5 and 6, MOSAIC produces a smaller *p*-value than Fisher’s method (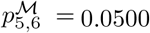 and *p*_5,6_ = 0.1998, respectively) and the other methods, indicating that MOSAIC is also a more powerful test for amplifying subgroup-specific mutual exclusivity patterns compared to *p*-value combination methods. These results illustrate that while MOSAIC loses some statistical power relative to subgroup-unaware approaches in the absence of subgroup-associated heterogeneity, it gains power when subgroup heterogeneity is present, outperforming standard *p*-value combination approaches in both settings.

### 3.2 Simulated Data

To show that MOSAIC is non-inferior to Fisher’s exact test for data without subgroup-dependent patterns of mutual exclusivity, we generated larger simulated datasets with numbers of patients that more closely resemble real-world genomics data. To do so, we repeatedly generated random matrices for 2 genes, 500 samples, and 2 patient subgroups using varying mutation frequencies and subgroup sizes. All mutations were generated independently of the subgroups. We evaluated each gene pair in each matrix with both MOSAIC and Fisher’s exact test. We used the Wilcoxon rank-sum test to compare the *p*-value distributions of both methods across 1,000 simulations of each unique combination of gene mutation rates and subgroup sizes; see Appendix A.2 for the parameters of the experiment.

The *p*-value distributions from MOSAIC and Fisher’s exact test did not significantly differ overall (median *p* = 0.59; Appendix Figures 1-3 and Appendix Tables 2-4) or on data with mutual exclusive mutations (median *p* = 0.54) according to Fisher’s exact test at a significance level of *α* = 0.05. These results indicate that MOSAIC, as a subgroup-aware method, produces comparable results to Fisher’s exact test, a subgroup-unaware method, on data with mutual exclusivity patterns unaffected by subgroup heterogeneity, demonstrating non-inferiority of our subgroup-aware approach.

### 3.3 Real-World Cancer Genomics Data

To show MOSAIC’s performance on real-world cancer genomics data, we sourced somatic mutation data and clinical metadata from The Cancer Genome Atlas (TCGA) Pan-Cancer Atlas [32]. First, we extracted somatic mutation data from the publicly-available MAF file and considered samples from primary solid tumors and the cancer type of interest [25]. We matched the samples to patient identifiers in the clinical metadata using the tumor sample barcode, only retaining missense mutations and samples with recorded clinical metadata for downstream analysis. Then, we created a binary mutation matrix described in Section 2. To restrict the number of gene sets for this analysis, we only considered genes that were mutated in at least 1% of all patients within a cancer type. Finally, we evaluated the candidate gene pairs using MOSAIC, as a subgroup-aware method, and Fisher’s exact test, a corresponding subgroup-unaware method. Given that multiple existing mutual exclusivity methods, including CoMEt and WeXT, reduce to Fisher’s exact test when evaluating pairs of genes, Fisher’s exact test serves as a representative, baseline approach for characterizing the impact of subgroup-awareness in mutual exclusivity analyses. Moreover, while CoMEt includes a subtype-aware formulation, we demonstrate that it does not differentiate between subgroup-associated and subgroup-unassociated mutual exclusivity (see Section 3.1), so we did not evaluate the subtype-aware formulation of CoMEt on real data.

The real-world cancer genomics analyses presented below serve as representative case studies, rather than an exhaustive benchmark. In particular, they highlight how our subgroup-aware approach distinguishes between different sources of mutual exclusivity in biological datasets and better contextualizes their interpretation in cancer biology.

#### Histological Subtype Analysis in Endometrial Cancer (UCEC)

Histology and genomics are inherently linked: the genomic profile of a tumor drives its phenotypic presentation. However, it remains unclear how underlying genomic variation between histological subtypes of a cancer type affects mutual exclusivity analyses. Endometrial cancer (UCEC) is comprised of multiple subtypes, including endometrioid endometrial adenocarcinoma (EEC) and serous endometrial adenocarcinoma (SEC) [34]. We stratified 527 UCEC patients into EEC (*n* = 394), SEC (*n* = 111), and a small mixed subgroup (*n* = 22).

In a genome-wide analysis of gene pairs in the UCEC cohort, MOSAIC’s histological subtype-aware analysis produces larger *p*-values for many gene pairs, relative to the subtype-unaware approach (Figure 3), with the largest relative differences in *p*-values reported for *TP53* and *KRAS, TP53* and *PTEN*, and *TP53* and *CTNNB1*. Inspection of the mutation frequencies of these known cancer driver genes reveals substantial differences between the SEC and EEC subtypes. *TP53* mutations are enriched in the SEC relative to the EEC, while *KRAS, PTEN*, and *CTNNB1* are enriched in the EEC relative to the SEC (Table 1). External studies also characterized *KRAS PTEN*, and *CTNNB1* mutations as distinct genomic features of EEC, and *TP53* mutations as a marker of SEC [21,29]. When both subtypes are evaluated in a pooled analysis, mutual exclusivity signals can arise that reflect the distinct genomic lineages of each subtype, rather than underlying biological interactions between genes. WeSME, a sophisticated, but subgroup-unaware analysis method, also identified *TP53, KRAS, PTEN*, and *CTNNB1* as a mutually exclusive gene set in UCEC [14]. MOSAIC, meanwhile, reduces this effect by controlling for variation between histological subtypes. Importantly, while these genes are known cancer driver genes, the patterns of mutual exclusivity between these genes are predominately caused by the different prevalences of mutations of these genes in different subtypes rather than the altered interactions between the genes.

**Fig. 3:**
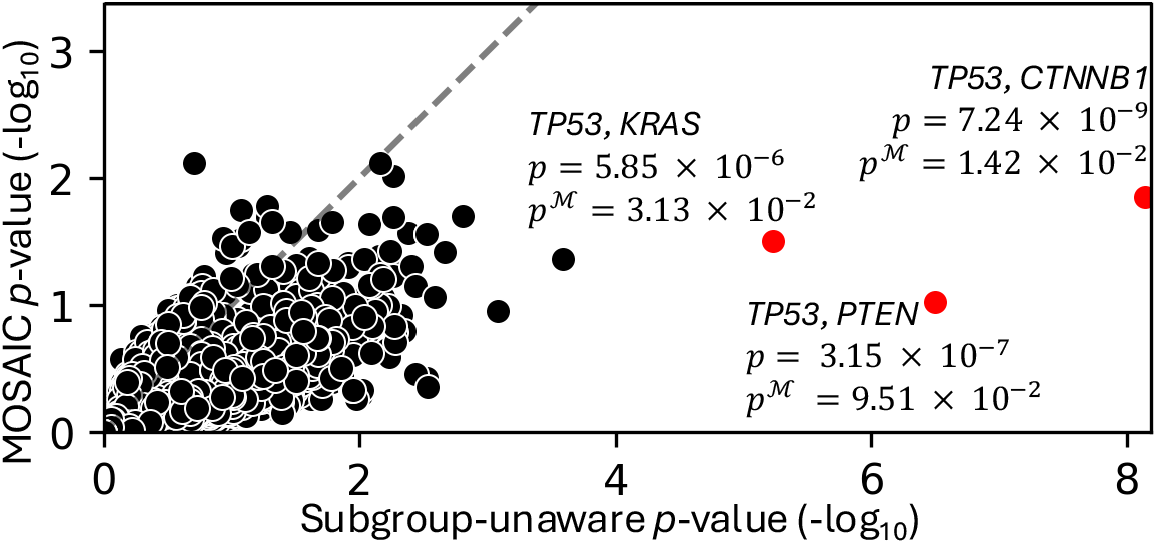
Comparison of the subgroup-unaware approach and MOSAIC’s histological subtype-aware approach in endometrial cancer (UCEC). Each point represents a gene pair evaluated by a subgroup-unaware method, Fisher’s exact test (*x*-axis) and our subgroup-aware method, MOSAIC (*y*-axis). Red points have the largest relative difference in *p*-values between the two approaches. The dashed line (*y* = *x*) indicates equal *p*-values.

**Table 1:** Histological subtypes of endometrial cancer (UCEC) have distinct genomic profiles. The mutation frequencies of *TP53, PTEN, KRAS*, and *CTNBB1* vary between the EEC and SEC subtypes of UCEC.

| | SEC ($n = 111$) | EEC ($n = 394$) |
| --- | --- | --- |
| *TP53* | 73.0% | 15.7% |
| *KRAS* | 3.6% | 24.1% |
| *PTEN* | 6.3% | 43.1% |
| *CTNNB1* | 0.9% | 32.5% |

#### Patient Age Group Analysis in Low Grade Glioma (LGG)

Older individuals typically accumulate more somatic mutations over their lifetimes and harbor higher mutational burdens than younger individuals [4]. WExT [18] controls for patient-specific mutation rates to reduce the number of false positive mutual exclusivity pairs introduced by variations in intertumor mutation rates. However, because this adjustment is conducted at the resolution of individual patients, these approaches cannot detect when a pattern of mutual exclusivity varies systematically across patient age groups. To assess how age-associated heterogeneity influences mutual exclusivity in low grade glioma (LGG), we split 508 patients into standard age subgroups: Adult (18-64 years old, *n* = 473) and Senior (65+ years old, *n* = 35). For simplicity, we excluded pediatric patients due to their small sample size (*n* = 2).

In the LGG cohort, *IDH1* and *IDH2* presented as the most statistically significant gene pair in the subgroup-unaware test (*p* = 2.61 × 10^*-*14^) and subgroup-aware test (*p*^ℳ^ = 3.98 × 10^*-*15^). However, inspection of *IDH1* and *IDH2* mutations in the cohort reveals that while both adult and senior patients exhibit high *IDH1* mutation rates, mutual exclusivity with *IDH2* mutations is found only in the adult subgroup (Figure 4). Moreover, although binning patients into Adult and Senior groups is a standard practice, the resulting subgroups were imbalanced and the adult subgroup may still contain age-associated variation. To address these limitations, we also stratified LGG patients into four groups based on quartiles of the cohort’s age distribution (Q1: 18–33 years, *n* = 144; Q2: 34–41 years, *n* = 118; Q3: 42–53 years, *n* = 123; Q4: 54–87 years, *n* = 123). With higher-resolution subgroups, MOSAIC localized the mutual exclusivity signal for *IDH1* and *IDH2* to younger patients (Q1–Q3, 18-53 years) with increased statistical significance (*p*^M^ = 5.55 × 10^*-*18^) compared to the analysis of Adult and Senior patients (*p*^ℳ^ = 3.98 × 10^*-*15^).

**Fig. 4:**
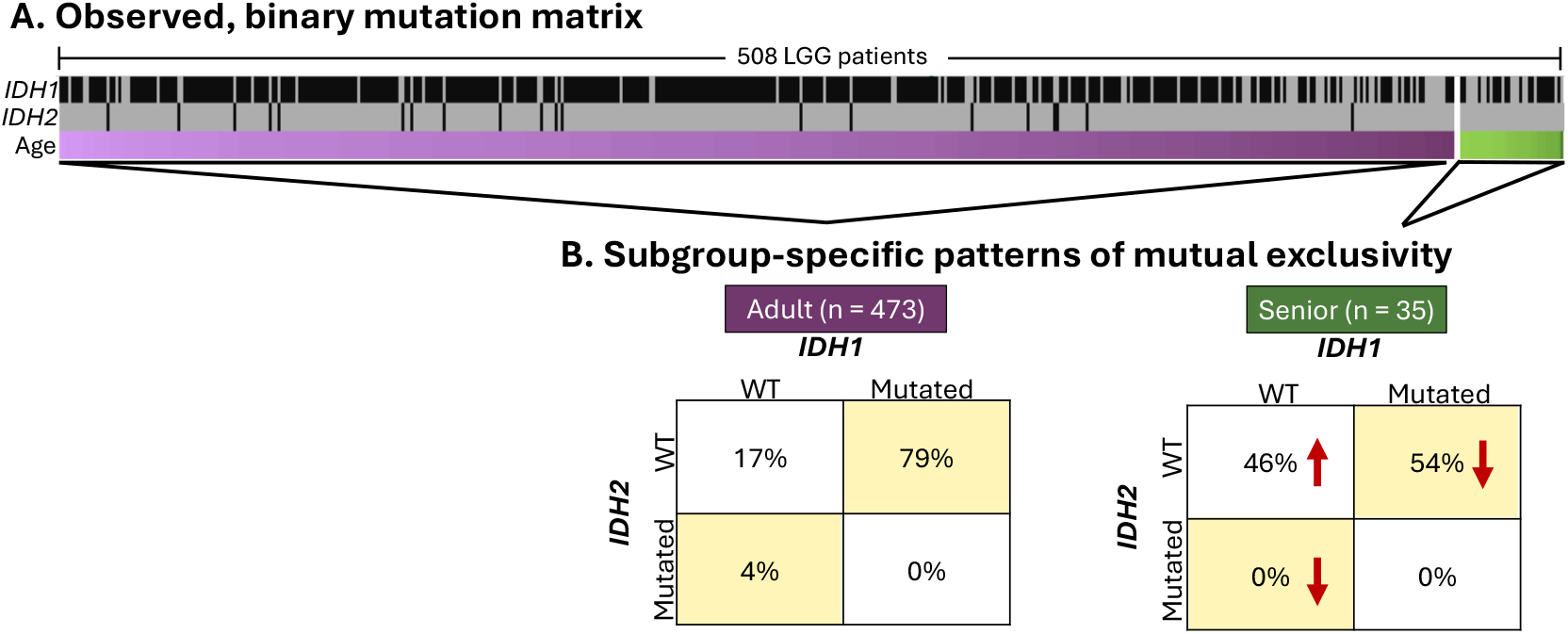
Age subgroup-aware characterization of *IDH1* and *IDH2* mutations in LGG. (A) Low-grade glioma (LGG) mutation data. Black bars indicate *IDH1* and *IDH2* mutations, top and middle rows. Color gradients indicate patient age (darker color denotes older patients in each age bin). (B) Contingency tables illustrating mutual exclusivity (yellow) in each age group. Red arrows denote mutation rate changes in Seniors (65+) relative to Adults (18-64).

Indeed, *IDH1* and *IDH2* present as statistically significant even in the subgroup-unaware test and have been externally validated as driver genes in LGG [10] [39], often associated with better prognosis due to the availability of *IDH* -targeted therapies [22]. Consistent with MOSAIC’s results, a meta-analysis of LGG genomic datasets reported that patients with *IDH* mutations are significantly younger than *IDH* -WT patients [38]. Thus, while *IDH1* and *IDH2* are known drivers in LGG, our analysis serves as a proof-of-concept that MOSAIC contextualizes the mutual exclusivity signal within biologically relevant subgroups.

## 4 Discussion

MOSAIC makes three key contributions. First, MOSAIC contextualizes mutual exclusivity signals that arise when the patient cohort is composed of distinct genomic lineages, as demonstrated with simulated data and in analyses of histological subtypes in endometrial cancer (UCEC). Second, in cases where mutual exclusivity is enriched in specific subgroups, as observed in simulated data and age-associated mutational patterns in low-grade glioma (LGG), MOSAIC localizes the subgroup-specific mutual exclusivity signal that would otherwise be diluted in pooled analyses. Finally, in datasets lacking biologically meaningful subgroup structure, MOSAIC is non-inferior to Fisher’s exact test. We also show that MOSAIC’s subgroup-aware approach is more powerful than Fisher’s method and other *post-hoc* meta analysis approaches for combining *p*-values across subgroups. Together, these results demonstrate that MOSAIC can help to distinguish between different sources of mutual exclusivity in genomics datasets.

This manuscript highlights how researchers may use MOSAIC to understand patient subgroups as a covariate in mutual exclusivity analyses. Users specify biologically-relevant subgroups *a priori*, according to available metadata for the sample cohort (e.g., clinical metadata for the TCGA cancer cohorts). As demonstrated in the low-grade glioma example, the test can be sensitive to sample subgroup definition. MOSAIC itself can be used to discover biologically relevant subgroups within a given cohort. Users may iteratively redefine patient subgroups to explore how a given subgroup definition impacts mutual exclusivity results relative to a subgroup-unaware approach. However, this alternative usage of MOSAIC is purely exploratory and should not be conducted on the same dataset used for downstream statistical inference. In practice, users who identify biologically relevant subgroups using MOSAIC must validate the significance of observed patterns of mutual exclusivity on external datasets.

For the simplicity of presentation, this manuscript does not explore all of the methodological details for MOSAIC, e.g., MOSAIC can support the mid *p*-value, which is less conservative than the approach described here [15]. Like other statistical methods, MOSAIC requires multiple testing correction for multiple gene sets. MOSAIC can restrict the evaluated gene sets *a priori* so as to not consider gene sets with insufficient numbers of mutations to survive multiple testing correction. Importantly, MOSAIC can be easily adapted to evaluate other signals in binarized data, such as co-occurring mutations.

In future work, MOSAIC will be extended to evaluate not just gene pairs, but larger gene sets. Additionally, while MOSAIC controls for gene mutation rates and patient subgroup heterogeneity, the statistical model should be refined to control for sample-specific mutation rates. Finally, due to the curse of dimensionality, MOSAIC is less efficient for *k* ≥ 4 subgroups. To account for this, we can implement an iterative computation with an early stopping condition. We begin the computation with the probability of observing the largest number of possible mutually exclusive mutations under the subgroup-aware null distribution and iteratively accumulate the tail probability, Pr(*X* ≥ *f*_*iℓ*_(*A*)), which monotonically increases by definition. The computation will complete when the cumulative probability exceeds a given threshold (i.e., 0.05) or the exact p-value is computed.

## 5 Conclusion

Mutual exclusivity tests can identify genes that may participate in shared biological pathways that, when functionally altered, drive tumorigenesis. However, existing methods cannot distinguish between mutual exclusivity patterns driven by underlying biological context from those influenced by patient subgroup heterogeneity. We introduced MOSAIC, a statistical model that controls for patient subgroup heterogeneity in mutual exclusivity analyses. As a result, MOSAIC improves the contextualization of candidate gene pairs and promotes more nuanced interpretation of mutual exclusivity signals in cancer.

## Acknowledgments

We thank Dr. Selen Bozkurt and Dr. Joyce Ho for their thoughtful feedback on this manuscript. We also thank Dr. Alexis Carter for her expert suggestions for the real-world genomics data preprocessing.

