## Appendix for "MOSAIC: Model-based, Subgroup-Aware Identification of Driver Mutations in Cancer"

### A Appendix

#### A.1 Comparison of MOSAIC with *p*-value combination methods for meta-analyses

| **Method** | ***p*-value for Gene 1 and 2** | ***p*-value for Gene 3 and 4** | ***p*-value for Gene 5 and 6** |
|---|---|---|---|
| MOSAIC | 0.0025 | 1.0000 | 0.0500 |
| Fisher's method | 0.0175 | 1.0000 | 0.1998 |
| Pearson's method | 0.0049 | 1.0000 | 1.0000 |
| Mudholkar & George's method | 0.0132 | 1.0000 | 1.0000 |
| Tippett's method | 0.0975 | 1.0000 | 0.0975 |
| Stouffer's method | 0.0100 | 1.0000 | 1.0000 |

Table 1: **Comparison of MOSAIC with *post-hoc* $p$-value combination methods.** For the toy dataset in Section 3.1, we evaluated three pairs of genes with two subgroups simultaneously using MOSAIC and independently using Fisher's exact test with several $p$-value combination methods.

#### A.2 Simulated Data Experiments

For these data, we generated random matrices with two genes with varying mutation rates (5%, 10%, 30%, and 50% of patients) and 500 patients with two subgroups of varying sizes (50% and 50%, 80% and 20%, and 90% and 10% of the patients). Since mutations were generated independently of the subgroups, any observed subgroup-associated patterns are due to chance alone. For each of the 48 configurations, we simulated 1,000 mutation matrices. For each configuration, we generated the $p$-value distributions resulting from the subgroup-unaware mutual exclusivity test, Fisher's exact test, and from MOSAIC's subgroup-aware approach. These were compared visually in histograms and quantitatively using the Wilcoxon rank sum test ($\alpha < 0.05$).

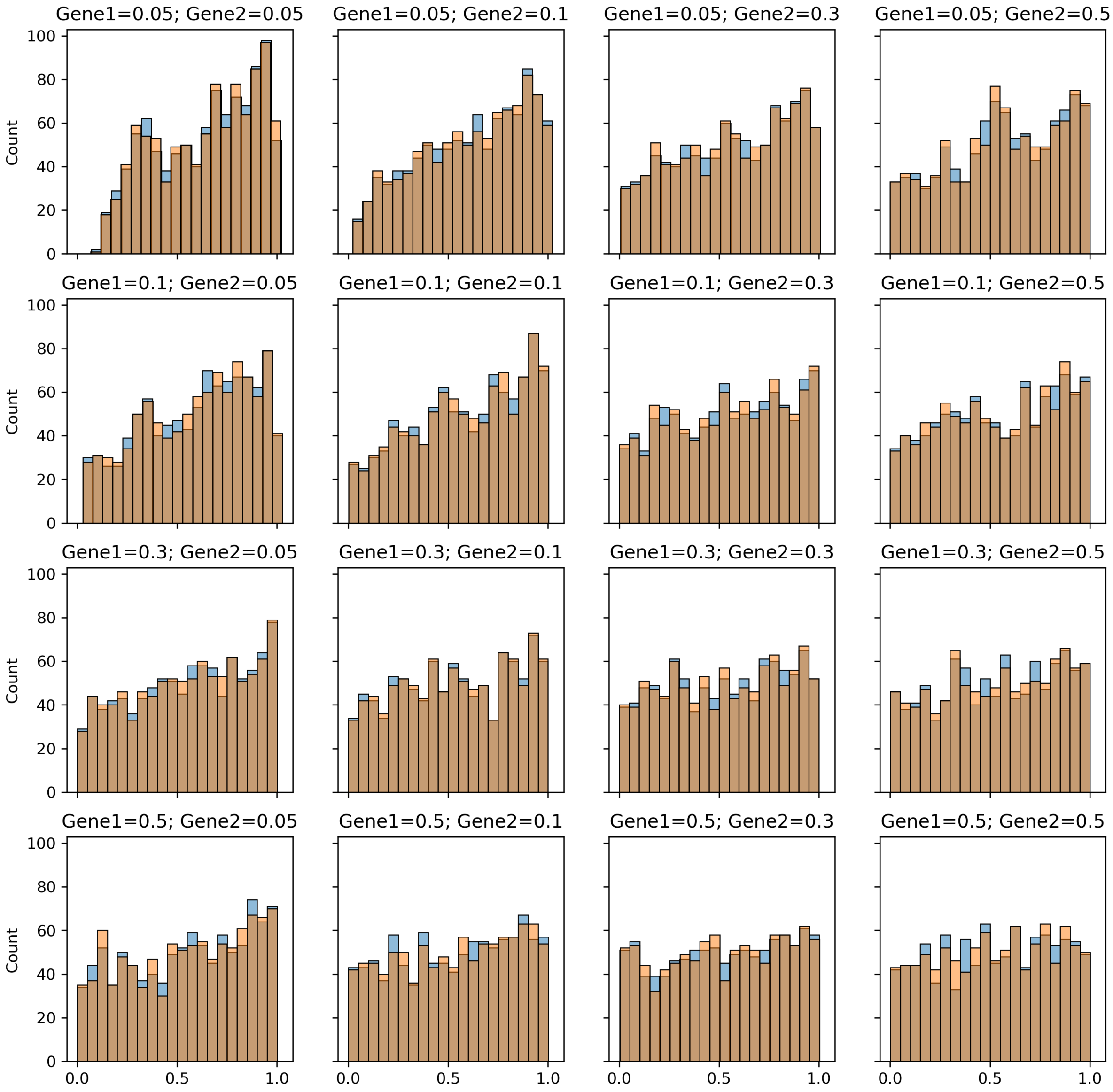


Fig. 1: Comparison of the resulting subgroup-unaware (orange) and subgroup-aware (blue) $p$-value distributions for simulated data. Each subplot title indicates the mutation rates for the two simulated genes. 500 patients were split into two balanced subgroups consisting of 250 patients each.

| | Gene 2: 0.05 | Gene 2: 0.1 | Gene 2: 0.3 | Gene 2: 0.5 |
|---|---|---|---|---|
| Gene 1: 0.05 | 0.8842 | 0.5128 | 0.1738 | 0.1746 |
| Gene 1: 0.1 | 0.1236 | 0.1449 | 0.8759 | 0.6253 |
| Gene 1: 0.3 | 0.8869 | 0.5361 | 0.7016 | 0.2697 |
| Gene 1: 0.5 | 0.6802 | 0.8636 | 0.8047 | 0.8240 |

Table 2: Wilcoxon rank-sum $p$-values comparing subgroup-aware and subgroup-unaware tests on simulated datasets without subgroup-associated variation. Each entry corresponds to a dataset defined by the mutation rates of gene 1 (rows) and gene 2 (columns). Simulations used two patient subgroups of size 250 and 250.

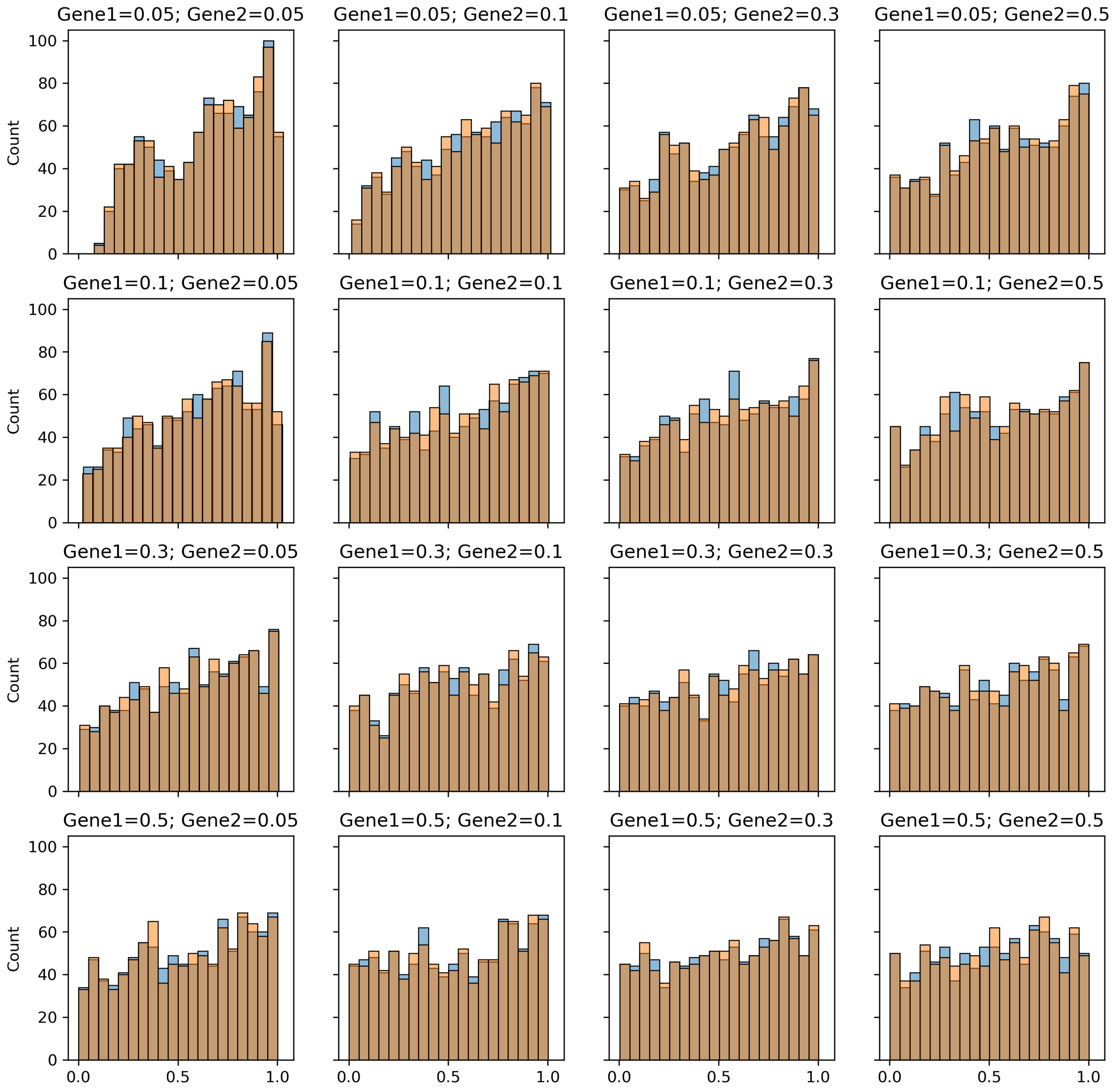


Fig. 2: Comparison of the resulting subgroup-unaware (orange) and subgroup-aware (blue) $p$-value distributions for simulated data. Each subplot title indicates the mutation rates for the two simulated genes. 500 patients were split into two imbalanced subgroups consisting of 400 and 100 patients, respectively.

| | Gene 2: 0.05 | Gene 2: 0.1 | Gene 2: 0.3 | Gene 2: 0.5 |
|---|---|---|---|---|
| Gene 1: 0.05 | 0.5892 | 0.8579 | 0.8770 | 0.3726 |
| Gene 1: 0.1 | 0.7711 | 0.3214 | 0.9655 | 0.4105 |
| Gene 1: 0.3 | 0.3813 | 0.9625 | 0.2629 | 0.5744 |
| Gene 1: 0.5 | 0.9493 | 0.4752 | 0.7870 | 0.4462 |

Table 3: Wilcoxon rank-sum $p$-values comparing subgroup-aware and subgroup-unaware tests on simulated datasets without subgroup-associated variation. Each entry corresponds to a dataset defined by the mutation rates of Gene 1 (rows) and Gene 2 (columns). Simulations used two patient subgroups of size 400 and 100.

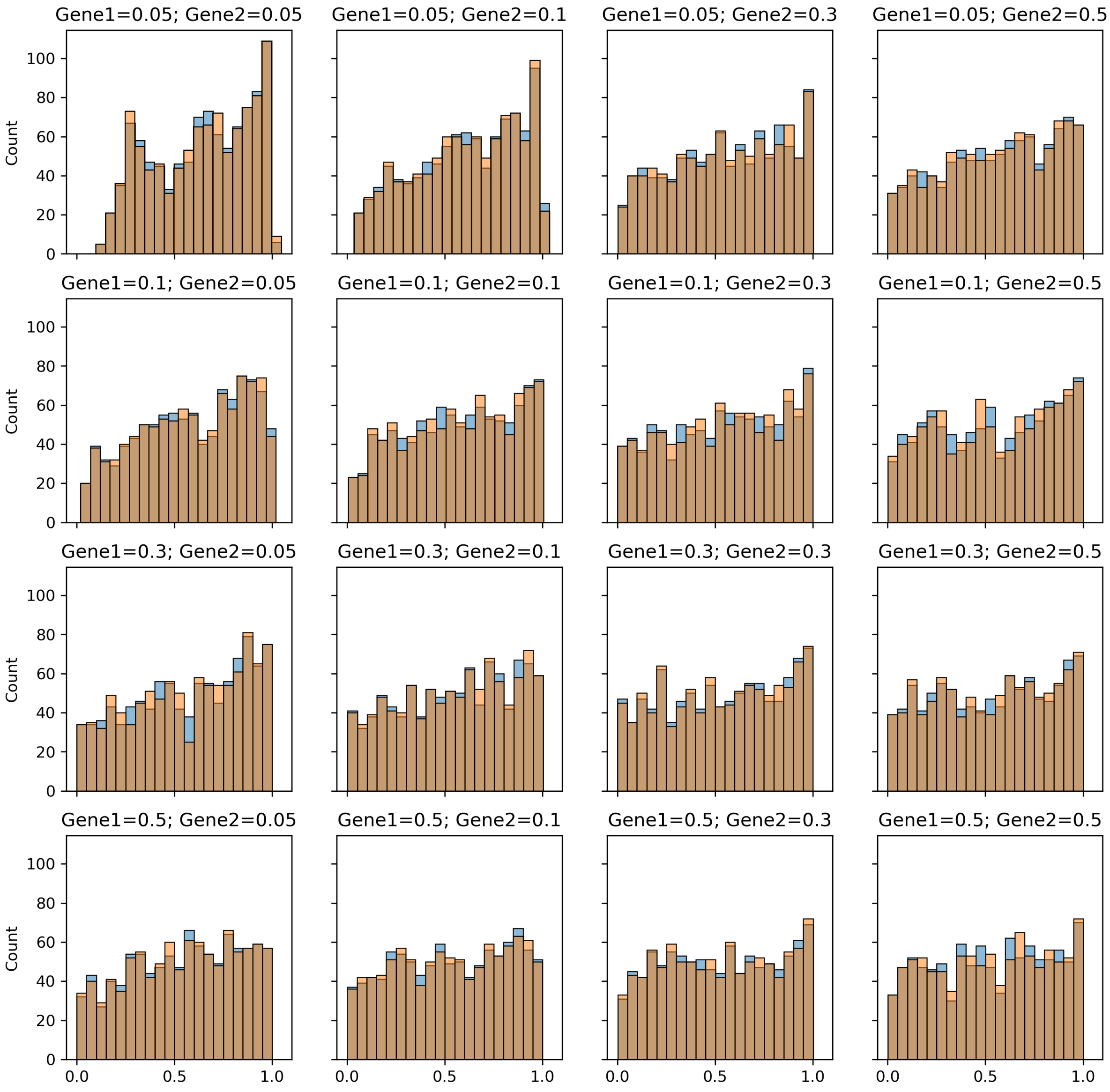


Fig. 3: Comparison of the resulting subgroup-unaware (orange) and subgroup-aware (blue) $p$-value distributions for simulated data. Each subplot title indicates the mutation rates for the two simulated genes. 500 patients were split into two highly imbalanced subgroups consisting of 450 and 50 patients, respectively.

| | Gene 2: 0.05 | Gene 2: 0.1 | Gene 2: 0.3 | Gene 2: 0.5 |
|---|---|---|---|---|
| Gene 1: 0.05 | 0.624399 | 0.762897 | 0.499780 | 0.840035 |
| Gene 1: 0.1 | 0.420133 | 0.147350 | 0.489680 | 0.538437 |
| Gene 1: 0.3 | 0.951383 | 0.822192 | 0.720223 | 0.364462 |
| Gene 1: 0.5 | 0.594432 | 0.374739 | 0.556952 | 0.634739 |

Table 4: Wilcoxon rank-sum $p$-values comparing subgroup-aware and subgroup-unaware tests on simulated datasets without subgroup-associated variation. Each entry corresponds to a dataset defined by the mutation rates of gene 1 (rows) and gene 2 (columns). Simulations used two patient subgroups of size 450 and 50.
